# Targeted single-cell genomics reveals novel host adaptation strategies of the symbiotic bacteria *Endozoicomonas* in *Acropora tenuis* coral

**DOI:** 10.1101/2022.04.22.489146

**Authors:** Keigo Ide, Yohei Nishikawa, Toru Maruyama, Yuko Tsukada, Masato Kogawa, Hiroki Takeda, Haruka Ito, Ryota Wagatsuma, Rimi Miyaoka, Yoshikatsu Nakano, Koji Kinjo, Michihiro Ito, Masahito Hosokawa, Kei Yura, Shoichiro Suda, Haruko Takeyama

## Abstract

*Endozoicomonas* bacteria symbiose with various marine organisms and are known to be beneficial for coral health. However, genome analysis of coral-associated *Endozoicomonas* has been limited owing to the difficulty in cultivation and metagenomic approach by contamination of host-derived sequences. In this study, we applied a novel single-cell genomics technique using droplet microfluidics to obtain single-cell amplified genome (SAGs) for coral-associated *Endozoicomonas* spp. genome. We obtained seven novel *Endozoicomonas* genomes from *Acropora tenuis* coral. These genomes revealed that *Endozoicomonas* bacteria played host-associated functions in host corals and had undergone independent host-adaptive evolution in different clades. These adaptive evolutions were mediated by host-derived eukaryotic-like genes, some of which were speculated to influence host immune mechanisms. These genes are speculated to enhance coral tolerance to environmental stresses. This study suggests the possibility of host adaptation of *Endozoicomonas* spp. in symbiosis with corals and their contribution to coral bleaching tolerance.

## Background

In recent years, coral bleaching has been known to have several impacts on marine ecosystems. Corals have symbionts, including intracellular symbiotic dinoflagellates, coral-associated bacteria, and viruses, which have been reported to play an important role in coral health and survival [1,2]. The importance of consideration for the function of the coral itself as well as the interactions among the holobionts in the coral ecology has been globally recognized. In the functional analysis of coral symbionts, symbiotic dinoflagellates have been the main target, but recently, the functions of coral-associated bacteria have also been gradually clarified, including the degradation of dead symbiotic dinoflagellates [3], provision of nitrogen sources [4], and protection from pathogenic bacteria via antibiotic activities [5].

The genus *Endozoicomonas* is the most pivotal coral-associated bacteria and is associated with an extensive range of marine organisms, including shellfish [6, 7], sea slugs [8], sponges [9, 10], and sea anemones [11, 12]. *Endozoicomonas* has been the most widely studied coral-associated bacterium, and it has been shown that phylogenetically different *Endozoicomonas* species colocalize within a single individual [3, 13]. In addition, *Endozoicomonas* has been reported to play several functional roles in coral survival, such as suppressing mitochondrial degradation, supplying sugars, and contributing to the coral sulfur cycle through dimethylsulphoniopropionate (DMSP) [14, 15]. However, little is known about the mechanisms by which *Endozoicomonas* establishes symbiotic relationships with its host corals. Recently, Ding J-Y et al. investigated the host cell entry mechanism of *Endozoicomonas montiporae* in *Montipora* coral, and their results suggested that this bacterium used eukaryotic-like genes, such as ephrin ligands, to enter the host cell [14]. This adaptation strategy is consistent with the fact that *Endozoicomonas* is not an obligate symbiont [14, 16]. However, considering its phylogenetic diversity, it is quite possible that *Endozoicomonas* employs other strategies. Moreover, cross-sectional analysis of genomic information of various *Endozoicomonas* species is necessary. Nevertheless, the number of draft genomes collected from the same host corals are limited, which hampers a comprehensive understanding of their symbiosis.

The lack of genomic information is due to the difficulty in culturing host-associated bacteria [16]. Thus, culture-independent approaches are needed for studying genome data to understand the relationship between *Endozoicomonas* and its host. Currently, shotgun metagenomics with binning is the mainstream method for microbial genome sequence analysis, but this method is greatly influenced by microbiome composition and diversity. In particular, in the genome analysis of closely related species with high similarity in 16S rRNA gene sequences, it is usually impossible to identify individual draft genome sequences by metagenome binning [17]. Furthermore, detailed genome sequence analysis of coral-associated bacteria is complicated because contamination by host mitochondria also significantly affects binning efficiency [16, 18]. As an alternative to metagenomics, single-cell genomics is very useful, as it can reveal the genome diversity of individual cells [19–21] from a mixed cell population.

Here, we suggest a novel approach for targeted single-cell genomics by detecting the target gene using droplet microfluidics. In this study, we newly identified seven draft genomes of *Endozoicomonas* bacteria from *Acropora tenuis* sampled at four different sites in Okinawa Prefecture, Japan. We then investigated the genomic differences between different clades of *Endozoicomonas* symbiotic with *A. tenuis* coral and examined the role of eukaryotic-like genes. Finally, we proposed a symbiotic strategy model for *Endozoicomonas* bacteria in *A. tenuis* corals, which revealed a diverse adaptation process.

The results of single-cell genome analysis indicate the existence of at least two strategies for microbe-host interactions, indicating the possibility of microbial regulation of host metabolic systems. These findings may provide insights into the robustness of coral reefs.

## Materials and Methods

### Coral branch sampling

*Acropora tenuis* samples were collected at four sites, namely Sesoko Minami (26°53.05’ N, 127°85.77’ E), Ishikawabaru (26°67.51’ N, 127°87.00’ E), Onna Village (26°50.36’ N, 127°85.39’ E), and the coral aquaculture center of Yomitan Village (26°40.90’ N, 127°71.55’ E) (Fig. 1A). Official permission to collect coral samples was obtained from Okinawa Prefecture in accordance with Okinawa Prefecture Fishing Regulation. The level of coral bleaching was evaluated on a 3-point scale (0: no bleaching, 1: slight bleaching, 2: complete bleaching). Small pieces of branches (~3 cm) were collected from *A. tenuis* corals and kept in a sterile zipper bag (140×100×0.04 mm; SEISANNIPPONSHA Ltd., Japan) filled with seawater from each sampling site. For 16S rRNA gene amplicon sequencing, coral branches were immediately transferred into a 5-mL tube (Eppendorf Ltd., Germany) containing 4 mL of RNAlater (Thermo Fisher Scientific, Waltham, MA, USA) and then freeze with liquid nitrogen. The frozen samples were kept in a −80°C deep freezer until analysis. For single-cell genome sequencing, the branches were kept on ice until just before the subsequent process.

**Figure 1.**
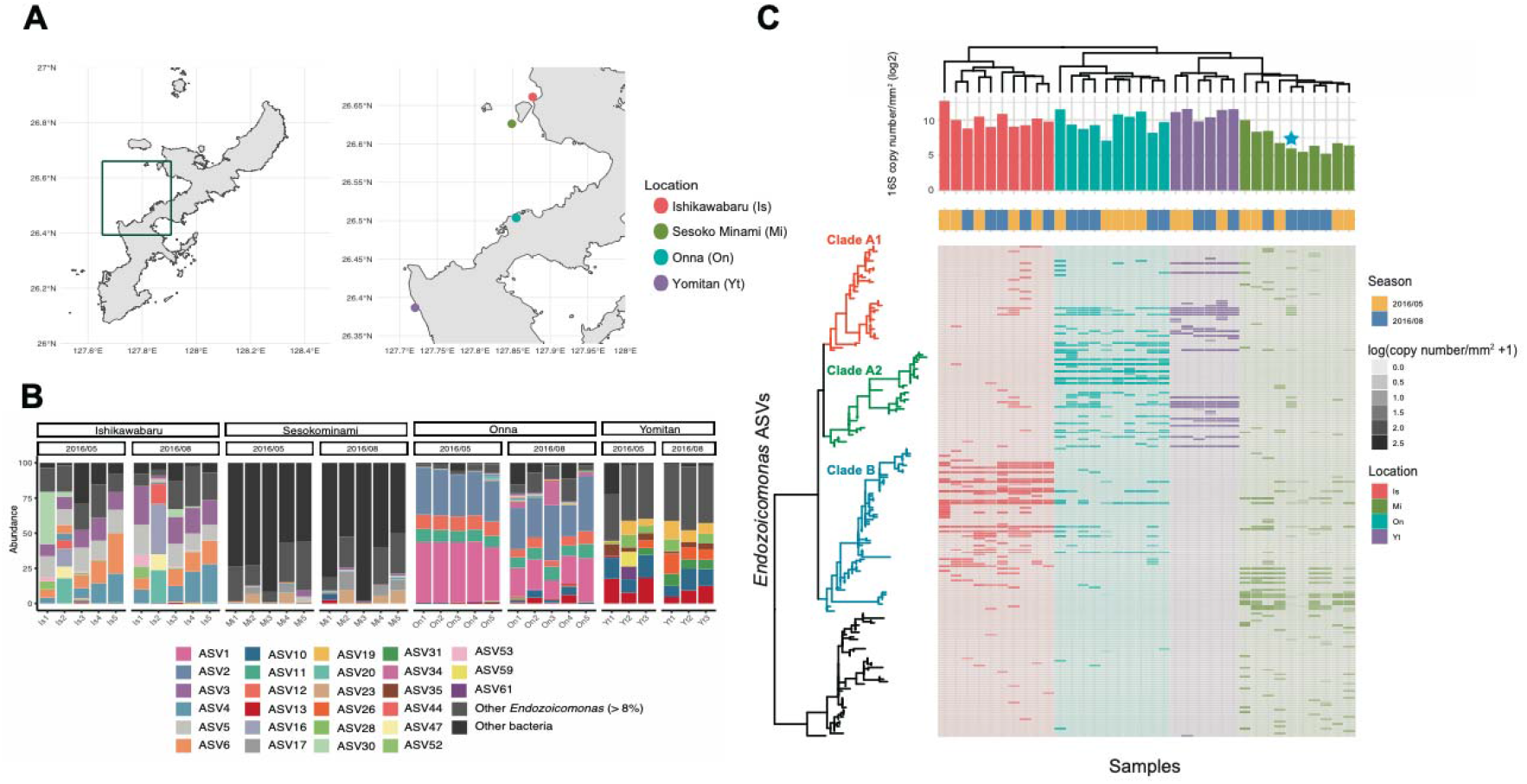
Distribution and diversity of the *Endozoicomonas* genus in *Acropora tenuis* from four sites in Japan. A: Coral samples were collected at four sites in Okinawa, Japan. Red: Ishikawabaru, Green: Sesoko Minami, Blue: Onna, Purple: Yomitan B: 16S rRNA gene composition. Each color indicates *Endozoicomonas,* and is shown for amplicon sequence variants that are above 8% in any sample. C: Heatmap plot based on 16S rRNA gene copy number denoting *Endozoicomonas* abundance. The copy number of 16S rRNA gene is normalized by the surface area of the coral. The uncultured Gammaproteobacteria clade (black) is displayed as the outer group. The star shows corals that did not bleach in August 2016. Bleaching was recorded in Ishikawabaru and Sesoko Minami, but not in Yomitan and Onna.

### DNA extraction and library preparation

The frozen coral branch was thawed on ice. The coral branch was then picked with tweezers and washed with artificial seawater. Then, in a sterilized zipper bag, the coral tissue was blown off using a Waterpik (EW-DJ61-W, Panasonic Corp., Japan), and the suspensions were collected into a 50-mL tube (Nippon Genetics Co., Ltd., Tokyo, Japan). After centrifugation at 10,000 × *g,* 4°C for 30 min, the supernatant was removed, and the pellet was resuspended in 500 μL of artificial seawater. The suspension was transferred into a 1.5-mL tube (SSIBio, Lodi, CA, USA) and centrifuged at 15,000 × g, 4°C for 15 min. After the supernatant was removed, the pellet was stored at −80°C until DNA extraction. DNeasy Plant Mini Kit (69104; QIAGEN, Hilden, Germany) was used for DNA extraction according to the manufacturer’s instruction, with a small modification. The thawed pellet was mixed with 400 μL of Buffer AP1 and transferred into 2-mL screw cap tubes with stabilized 0.1 mm zirconia beads (Yasui Kikai Corp., Osaka, Japan). The tubes were homogenized three times at 2,500 rpm for 60 s at 60-s intervals. After centrifugation, 4 μL of RNase A were added, and the sample was incubated at 65°C for 10 min before DNA extraction was performed following the kit protocol.

### 16S rRNA gene sequencing and analysis

The bacterial community was identified by targeting the variable region V1-V2 of the 16S rRNA gene using the primer set 27F and 338R. PCR amplification of the 16S rRNA gene amplicon (300 bp) was performed in a 25-μL mixture, and the amplicons were sequenced using Ion Torrent PGM with 318 Chip v2 (Thermo Fisher Scientific).

16S rRNA gene analysis was performed using QIIME2 2020.2 [22]. For quality control, amplicon sequence variants (ASVs) were constructed using DADA2 (via q2 dada2) [23]. All ASVs were aligned using mafft (via q2-alignment) [24]. Phylogenetic tree was constructed based on masked aligned ASVs using fasttree2 (via q2-mask and q2-phylogeny) [25]. Taxonomic assignment was performed using the q2-feature-classifier [26] classify-sklearn naiūve Bayes taxonomy classifier against the Silva 138 99% OTUs full-length sequences [27]. The downstream analysis was performed using the R bioconductor phyloseq [28] and ggtree [29] in R version 3.5.2.

### Droplet digital PCR for quantification of 16S rRNA gene copy number

The PCR mixture contained 10 μL of 2× ddPCR Supermix for Probes (Bio-Rad, Hercules, CA, USA), 1.0 μL of 10 μM forward and reverse primers, 1.0 μL of 10 μM Taqman probe, 1.0 μL of 1 mM Dextran fluorescein (FD10S; Merck KGaA, Darmstadt, Germany), 1.0 μL of extracted DNA, and 5.0 μL of nuclease-free water (Sigma-Aldrich, St. Louis, Missouri, USA). A total of 20 μL of the PCR mixture were introduced into microfluidic devices (Supplemental figure 1), and 40 μM of droplets were generated with droplet generation oil for probes (Bio-Rad). The primers and Taqman probes were synthesized by Integrated DNA Technologies (IDT, Inc., Coralville, IA). The sequence of the forward primer, reverse primer, and Taqman probe was 5’-AGA GTT TGA TCM TGG CTC AG-3’, 5’-GCT GCC TCC CGT AGG AGT-3’, and [6-FAM]-5’-CAG GCC TAA-[ZEN]-CAC ATG CAA GTC-[IBFQ]-3’, respectively. The droplets were collected into 0.2-mL tubes and then thermal cycled using a T100™ Thermal Cycler (Bio-Rad) with the following protocol: 95°C for 10 min, followed by 40 cycles of 94°C for 30 s and 60°C for 1 min, 95°C for 10 min, and maintenance at 4°C. Subsequently, the droplets were reintroduced into microfluidic devices for digital counting. We have developed a system for calculating the rate of fluorescence-positive droplets using a 488-nm laser (Lambda mini, Tokyo Instruments, Japan) (Supplementary Fig. S1). Fluorescent signals from the droplets were detected by a photosensor module (H11902-01; Hamamatsu Photonics KK, Hamamatsu, Japan) and recorded by an oscilloscope (PICOSCOPE 2204A; Pico Technology Ltd., St. Neots, Cambridgeshire, UK). Combined with the information of droplet diameter measured with ImageJ, 16S rRNA gene copy number per 1 ng of extracted DNA was calculated [30].

### Measurement of surface area of coral skeletons by CT scan

The coral branches with surface tissue removed by water picking were used for calculating the surface area. 3D images of the coral branches were obtained using an X-Ray CT scanner (TDM1300-IS; Yamato Scientific, Japan). The surface area was calculated by the Mimics software (Materialise, Leuven, Belgium). The 16S rRNA gene copy number per 1 cm^2^ of surface area was calculated.

### Preparation of bacterial suspension for single-cell genome sequencing

A small piece of coral branch was collected and kept in a 25-mL tube (Iwaki, Tokyo, Japan) filled with seawater from each sampling site. The tube was kept on ice immediately. Each branch was transferred into 5 mL of 0.22-μL-filtered UV-treated seawater of each sampling site and crushed thoroughly by a scalpel (Feather Safety Razor Co. Ltd., Japan).

After 5 min of standing on ice, the supernatant was collected into three 1.5-mL tubes. The tube was subsequently centrifuged at 300 × *g* for 5 min, and the supernatant was transferred into a new 1.5-mL tube to remove symbiotic dinoflagellates and other larger particles. The collected supernatant was centrifuged at 8,000 × *g* for 5 min. The supernatant was removed, and 500 μL of 0.22-μL-filtered UV-treated seawater was added to resuspend the pellet. The centrifugation at 300 × *g* was repeated until most of the symbiotic dinoflagellates were removed. The centrifugation at 8,000 × *g* and the washing steps were repeated three times.

Next, the pellet was resuspended in 500 μL of 1x SYBR Green (Thermo Fisher Scientific, Hampton, NH) for 5 min for DNA staining. After centrifugation at 8,000 × *g,* the pellet was resuspended in 50 μL of 0.22-μL-filtered UV-treated seawater, and the cell concentration was calculated with a bacterial counter (SLGC, Tokyo, Japan) under a fluorescent microscope (CKX53; Olympus Optical Co Ltd., Tokyo, Japan).

### Whole genome amplification and detection of *Endozoicomonas* in gel beads

Cells were encapsulated into 40-μm microfluidic droplets at a concentration of 0.3 cell/droplet using a microfabricated microfluidic device reported previously [19]. The theoretical number of encapsulated cells in a droplet was derived by the Poisson distribution (Formula 1). The percentage of doublets would be less than 3.7% of the total droplets when the cell concentration was 0.3 cell/droplet.

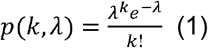

Next, gel-beads based whole genome amplification (WGA) was conducted according to the previously reported WGA method (single-cell amplified genomes (SAGs) in the gel are referred to as SAG-gel) [21]. After WGA, the gel beads were stained with DAPI, and amplification of the DNA was confirmed with microscopic observation (CKX53; OLYMPUS). Subsequently, the gel beads were washed and suspended in UV-treated Dulbecco’s phosphate-buffered saline (-) (Thermo Fisher Scientific). To detect the target gene sequence, 20 μL of PCR mixture consisting of 10 μL of PrimeTime Gene Expression Master Mix (Integrated DNA Technologies, Coralville, IA, USA), 1.0 μL of each 10 μM primer (forward and reverse, final concentration 500 nM), 0.5 μL of 10 μM Taqman probe (final concentration 250 nM), and 7.5 μL of gel beads solution was prepared. The sequence of each primer and probe is listed in Supplementary Table 1. On the basis of the 16S rRNA gene amplicon sequencing results, we designed two sets of primers and probes (Sets A and B) using the Geneious software (Biomatters Ltd., Auckland, New Zealand). The primer and probe sequences were evaluated with Primer BLAST [31] to confirm that they did not exhibit any specificity to the mitochondrial genome of *A. tenuis.* To detect the *Endozoicomonas* sequence amplified in gel beads, either Set A or Set B was used. The PCR mixture was reintroduced into microfluidic devices, and 50 μm of gel beads-containing microfluidic droplets were generated with carrier oil (HFE-7500 oil with 2% surfactant; RAN Biotechnologies Inc., Beverly, MA, USA). The concentration of the beads was adjusted at 0.3 bead/droplet. The droplets were collected into a 0.2-mL tube, and PCR amplification (95°C for 3 min, 25 cycles of 95°C for 5 s and 50°C for 40 s, and maintenance at 10°C) was conducted. After thermal cycling, the fluorescence of the droplets was observed with a microscope (CKX53; OLYMPUS). The droplets exhibiting both DAPI and FAM fluorescence were manually picked with a micro dispenser (Drummond Science Company Broomall, PA, USA) and transferred into a 0.2-mL tube. Next, second-round multiple displacement amplification (MDA) was performed using a REPLI-g Single Cell Kit (Thermo Fisher Scientific). Buffer D2 (0.6 μL) was added to each well, followed by incubation at 65°C for 10 min. After that, 8.6 μL of MDA mixture (0.6 μL of stop solution, 1.8 μL of H_2_O, 5.8 μL of reaction buffer, and 0.4 μL of DNA polymerase) was added, and the mixture was incubated at 30°C for 120 min. The MDA reaction was terminated by heating at 65°C for 3 min. The amplicon yields were quantified by a Qubit dsDNA HS assay kit (Thermo Fisher Scientific). PCR amplification over the 16S rRNA gene was also performed against second-round MDA products. Primer pair sequences for the V3-V4 region were used according to Illumina’s

MiSeq system protocols (Forward: 5’-TCG TCG GCA GCG TCA GAT GTG TAT AAG AGA CAG CCT ACG GGN GGC WGC AG-3’, Reverse: 5’-GTC TCG TGG GCT CGG AGA TGT GTA TAA GAG ACA GGA CTA CHV GGG TAT CTA ATC C-3’). PCR amplification was confirmed with agarose electrophoresis (100 V, 15 min), and the amplicon sequences were obtained via Sanger sequencing (Fasmac, Kanagawa, JAPAN) to determine whether the target *Endozoicomonas* sequence was acquired.

### *Endozoicomonas* single-cell genome sequencing and genome assembly

Second-round MDA products from single-cell samples were used for next-generation sequencing library preparation with Nextera XT DNA sample prep kit (Illumina) according to the manufacturer’s instructions. Each SAG library was sequenced using an Illumina MiSeq 2 × 75bp.

All raw reads were trimmed to remove adapter sequences by fastp v0.20.0 with default parameters [32]. Trimmed reads were assembled using spades v3.12.0 with ‘-k auto --sc --careful’ parameters [33]. All SAGs were concatenated using ccSAG [34]. SAGs with an average similarity of > 99.9% in single-copy marker genes were co-assembled and used for subsequent analyses. Contigs less than 1,000 bp in the SAG were removed. We removed host- and human-derived contaminations, which were annotated as eukaryota by BLAST against the NCBI nt database, from the SAG [35]. The quality of the SAGs was checked using CheckM v1.0.13 [36]. For the subsequent analyses, only high-quality SAG data were used (85% > completeness, < 5% contamination).

### Phylogenetic and comparative genome analysis

Gene prediction was performed to estimate protein coding genes (prodigal v. 2.6.3) [37], tRNAs (aragorn v. 1.2.38) [38], and rRNAs (Barrnap v. 0.9) [39] by Prokka v.1.14.5 for the SAGs [40]. Predicted proteins were annotated by diamond v0.9.14 [41] against the NCBI nr database [42]. We also ran InterProScan v.5.40-77.0 [43] and eggnog-mapper v.1.0.3 [44] to annotate domain information and GO terms and KEGG orthology. Enrichment analysis was performed using the clusterProfiler package [45].

The orthogroup was estimated using Orthofinder v2.3.0 with the default parameters [46]. Single-copy marker genes were extracted based on the Orthofinder results. Each gene was aligned by mafft v7.407 and trimmed by trimal v1.4 [24, 47]. A phylogenetic tree was constructed using the single-copy marker genes by IQ-TREE v1.6.7 [48] with ultrafast bootstrap [49] and ModelFinder [50]. Average Nucleotide Identity (ANI) and Average Amino Acid Identity (AAI) were calculated by the All against all ANI/AAI matrix calculator [51].

### Eukaryotic-like protein analysis

All the Pfam IDs with a Z-score of 10000 in EffectiveELD version 5.2 were extracted from the Interproscan results [52]. For extracting eukaryotic-like genes based on sequence homology, genes were selected by taxonomic annotation using eggnog-mapper v.1.0.3 [44]. In addition, we performed a taxonomy annotation of eukaryotic-like genes using mmseqs2 with “--tax-lineage 1 --vote-mode 0” options, using the eukaryotic-only NCBI nr database. We also checked that these genes are not annotated with other bacteria to confirm that *Endozoicomonas* acquired the genes from eukaryotes during host adaptation [53] by using the NCBI nr database without the *Endozoicomonas* genus. Major hits of annotation result were segmented annelid worms *Capitella teleta.* These genes were considered to be derived from contamination and removed from the results.

## Results

### Diversity analysis of the *Endozoicomonas* genus in *Acropora tenuis*

*A. tenuis* were collected at four locations (Ishikawabaru, Sesoko Minami, Yomitan, and Onna) in Okinawa in May and August 2016 (Figure. 1A). These four locations are placed within a 15-km radius. Through 16S rRNA gene sequencing, 1,418 ASVs were detected as coral-associated bacteria, and 209 ASVs were classified as *Endozoicomonas* spp. In Ishikawabaru, Yomitan, and Onna, *Endozoicomonas* spp. was dominant, representing 70.7-99.7% of the detected bacteria (Figure. 1B). However, in Sesoko Minami, the proportion of *Endozoicomonas* spp. was only 1.8-50.3%, and order Rickettsiales was dominant instead (Supplementary Figure 2A), including Rickettsiales bacteria strain SESOKO1, which was previously reported by our group [54]. The results of Principal Coordinate Analysis (PCoA) based on the Bray Curtis distance of *Endozoicomonas* composition showed spatial specificity for *A. tenuis* (Supplementary Figure 2B and 2C). Phylogenetic analysis of the 16S rRNA gene revealed that the *Endozoicomonas* species can be divided into three major clades: Clade A1 (Red) and Clade A2 (Green) constituted the Onna and Yomitan samples, whereas Clade B (Blue) comprised the Ishikawabaru and Sesoko Minami samples (Figure 1C, phylogenetic tree and heatmap). Absolute abundance of 16S rRNA gene was calculated by the 16S rRNA gene copy number measured by ddPCR and the surface area value obtained by CT scan. The average (±sd) number of *Endozoicomonas* was 37952 ± 75836 copies/mm^2^ (Ishikawabaru), 2306.8 ± 4241.6 copies/mm^2^ (Sesoko Minami), 30180 ± 30165 copies/mm^2^ (Onna), and 37287 ± 29083 copies/mm^2^ (Yomitan), respectively (Figure 1C, bar plot). These results suggest that the cell numbers of *Endozoicomonas* spp. vary drastically from site to site. Especially at Sesoko Minami, where the copy number of 16S rRNA gene was exceedingly low, the absolute number of *Endozoicomonas* was significantly lower than that at other sites. In August 2016, large-scale coral bleaching was observed around Okinawa Prefecture; however, the bleaching level of coral colonies at Sesoko Minami was lower than that at Ishikawabaru [55]. In particular, the single coral colony, in which *Endozoicomonas* Clade A1 was dominant, did not show any bleaching in August 2016 (Figure 1C; star). In our previous study, the density per surface area of zooxanthellae in corals was greatly reduced during coral bleaching in August 2016 [55]. However, in the same coral, the absolute number of *Endozoicomonas* did not decrease significantly from May to August in 2016 (Figure 1C).

### Single-cell genome sequencing of *Endozoicomonas* bacteria

To selectively obtain genomic information of target bacteria from crushed coral tissue suspensions, we have developed a method for targeted single cell genomics using droplet microfluidics. Bacterial suspensions prepared from coral tissues were encapsulated into agarose gel droplets and processed to gel-beads based WGA to obtain SAG-gel [21] (Fig. 2a). Microscopic observation revealed that 24.4-26.3% of the gel beads exhibited DAPI fluorescence, which corresponded to the Poisson distribution value. Subsequently, PCR-based detection of target genes with FAM probes was conducted, which enabled the identification of gel beads containing target SAGs (Fig. 2b). After PCR amplification, there were droplets displaying both DAPI and FAM fluorescence (Fig. 2c). The percentage of DAPI fluorescence-positive droplets is 0.8-1.2%, which represents the rate of droplets encapsulating the target bacterium.

**Figure 2.**
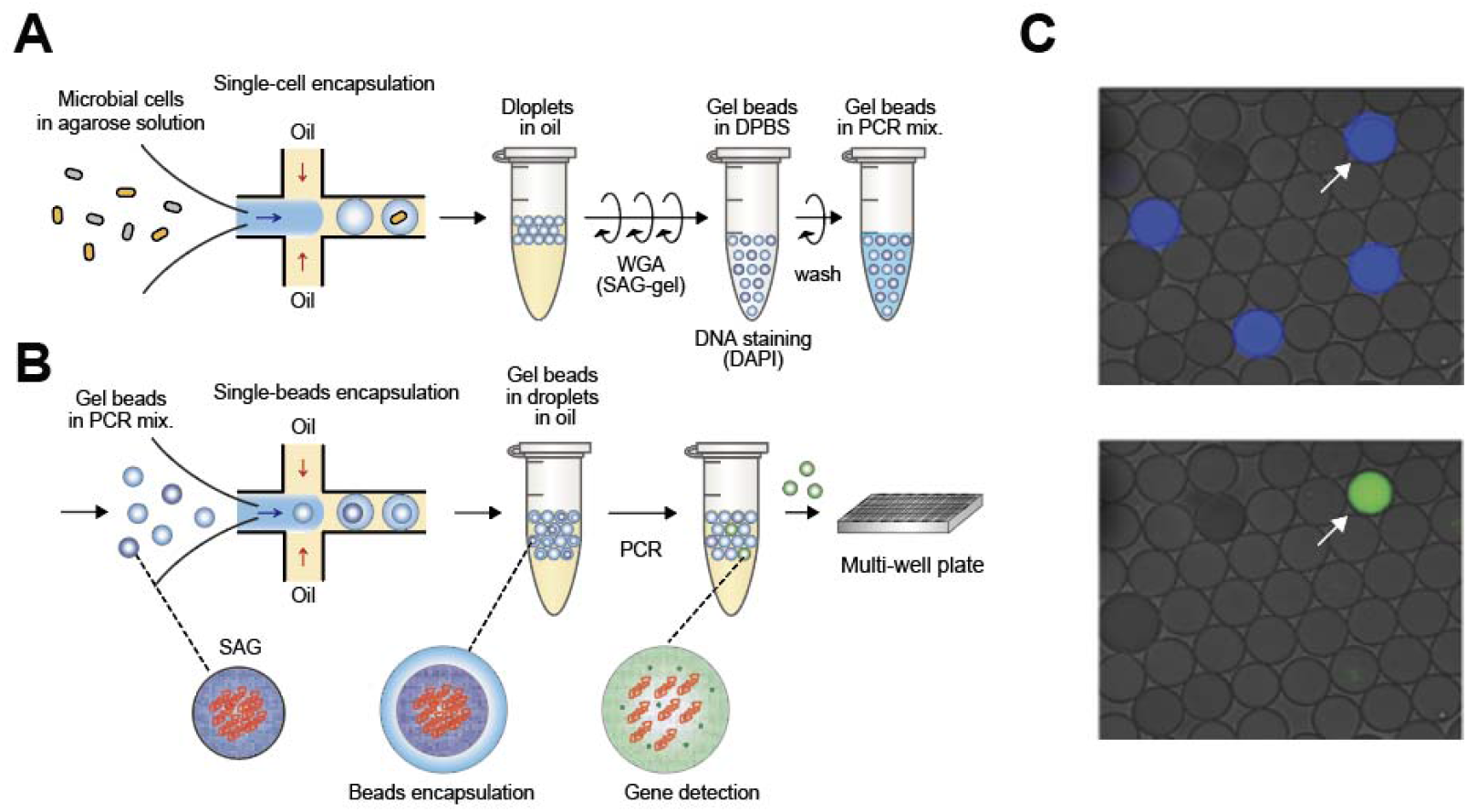
Targeted single-cell genomics with droplet microfluidics. A: Microbial cells were encapsulated into 40-μm microfluidic droplets at the single-cell level (0.3 cell/droplet) with ultra-low melting temperature agarose solutions. Collected droplets were solidified, treated with lysis solution, and subjected to whole genome amplification (WGA) using a previously described single-cell amplified genome (SAG)-gel platform. After washing, whole genome amplified gel beads were resuspended in the PCR mixture. B: Gel beads suspended in PCR mixture were reintroduced into microfluidic devices, and Water in Oil (W/O) droplets encapsulating gel beads were generated. After thermal cycling, droplets with target sequence exhibited fluorescence derived from Taqman probes. The fluorescence-positive droplets were isolated into multi-well plates and then subjected to second-round WGA and library preparation. C: Microscopic images of W/O droplets after PCR. Droplets with genome amplification by MDA exhibit blue fluorescence derived from DAPI. Droplets with target gene amplification by PCR exhibit green fluorescence derived from FAM. The droplet indicated by the arrow head contains genome amplicons of the target bacterium.

The results of 16S rRNA gene sequencing followed by second-round MDA showed that 64% (28 out of 44) of the samples corresponded to the *Endozoicomonas* sequence. Among the remaining 16 samples, 13 exhibited insufficient DNA yield (< 2 ng/μL) after the second-round MDA. The remaining three were confirmed to be derived from host mitochondria. When droplets that were positive only for DAPI were isolated, all the sequence corresponded to the sequence of host mitochondria (n=30), suggesting that our targeted single-cell genomics has a low false-negative rate. In order to obtain 28 target bacterial SAGs by random sampling, 2333-3500 droplets need to be isolated according to the calculation, which means that our method showed 53.0-79.5 times higher screening efficiency than the random sampling.

As a result, seven draft SAGs ranging from 2.1 Mbp to 6.4 Mbp were acquired from four sampling sites, three of which were classified as high-quality and three of which were classified as medium-quality according to the criteria by the Genomic Standards Consortium (GSC) (Table. 1).

### Comparative genome analysis of *Endozoicomonas* genus

We conducted a comparative genome analysis using our five draft SAGs, which include four from this study and one from our previous report [55], collected from *A. tenuis,* and 10 public *Endozoicomonas* SAGs collected from various kinds of marine organisms. Phylogenetic trees constructed with single-copy marker genes showed that *Endozoicomonas* spp. represented single clades that corresponded to symbiotic host organisms. However, *Endozoicomonas* spp. collected from *Acropora* coral *(Endozoicomonas* sp. ISHI1, SESOKO1, ONNA1, ONNA2, and YOMITAN1) were divided into two clades (Clade A1 and B) (Figure 3A), which was also confirmed by the ANI and average AAI (Supplementary Figure 3).

**Figure 3.**
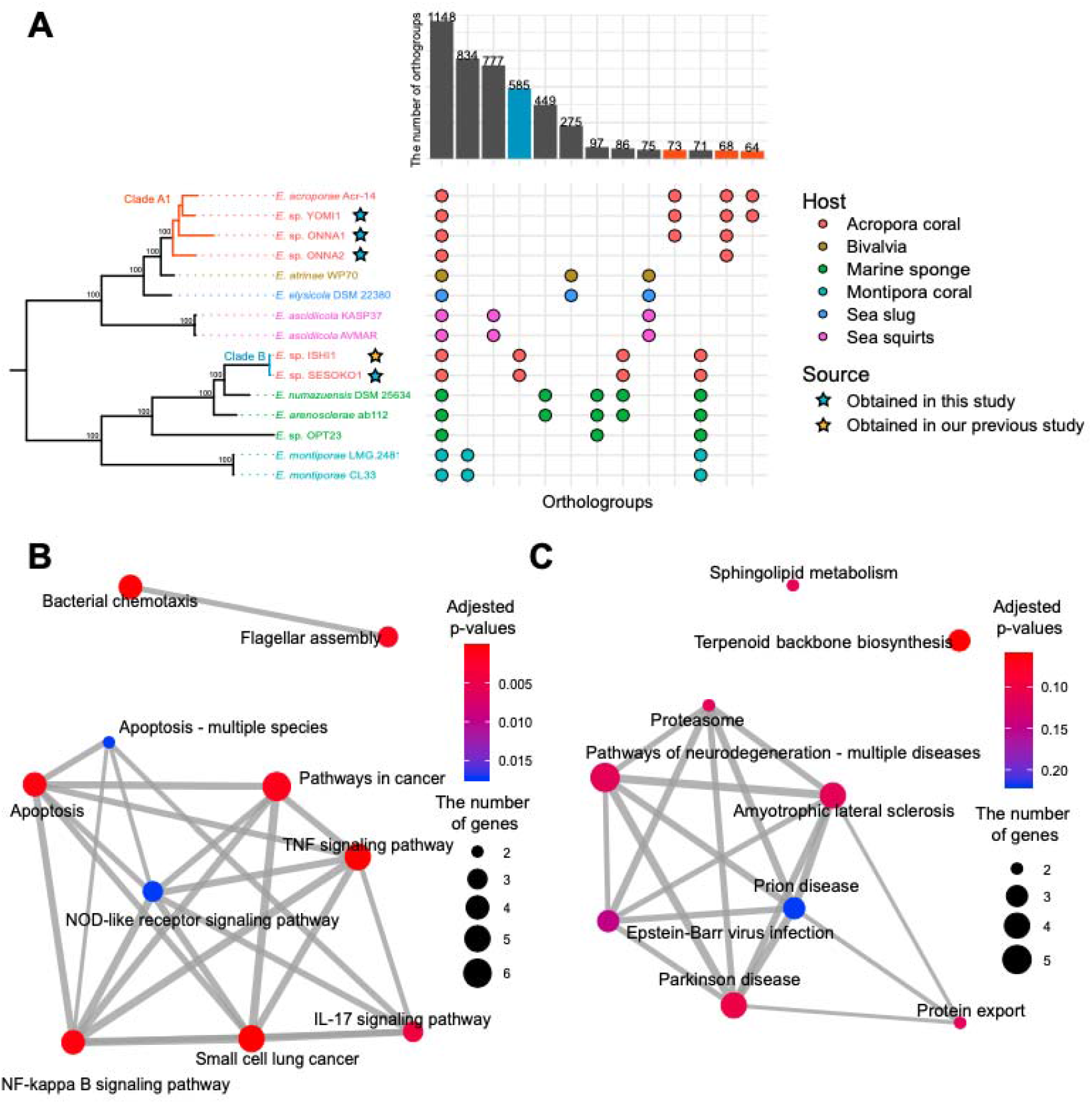
Comparative genome analysis of *Endozoicomonas*. A: Plot of lineage-specific genes based on ortholog analysis. Bar plots indicate the number of orthogroup types. Clade A1 and Clade B correspond to the clades in Fig. 1C based on the similarity of the 16S rRNA genes. The color of the circles and species names indicate the type of host organism. The colors of the bar plot indicate the genes used in the KEGG enrichment analysis shown in B (red: clade A1 specific) and C (blue: clade B specific). B: Enrichment analysis results of *Endozoicomonas* Clade A1-specific genes for KEGG pathways. The top 10 pathways are shown. The size of the circle indicates the number of genes. The thickness of the lines between pathways indicates the number of shared genes. C: Enrichment analysis results of *Endozoicomonas* Clade B-specific genes for KEGG pathways.

A similar trend was observed in the ortholog analysis. Out of a total of 17475 orthogroups, the number of orthogroups commonly detected in all *Endozoicomonas* spp. was 1,148. We confirmed that the bacterial secretion systems are evolutionarily conserved in all *Endozoicomonas* symbiotic relationships with marine organisms (Supplementary Table 2), which are important for *Endozoicomonas* to secrete proteins related to eukaryotic pathways in the host. The prediction of effector proteins showed that 12.5-20.4% (T3SS) and 2.97-4.22% (T4SS) of proteins were released by bacterial secretion systems (Supplementary Table 3). On the other hand, the remaining 16,327 orthogroups were clade-or strain-specific, suggesting that *Endozoicomonas* spp. possess a diverse orthogroup profile. It appears that the profile of the orthogroups depends on the host of each *Endozoicomonas* sp. However, Clade A1 and Clade B, which were symbiotic with the same host (*A. tenuis),* shared no unique orthogroups.

Enrichment analysis of accessory genes to the KEGG pathway revealed the functional characteristics and differences of each clade (Clade A1 and Clade B). Of the top ten enriched pathways (nine pathway in Clade B) of each clade, eight pathways (Except Bacterial chemotaxis and Flagellar assembly) in Clade A1 as well as seven pathway (Except Sphingolipid metabolism and Terpenoid backbone biosynthesis) in Clade B were assigned as eukaryotic pathways (Figure 3B,C). Primarily, in *Endozoicomonas* Clade A1, immune-related pathways were enriched, such as the TRAF signaling pathways and NF-κB pathways (Figure 3B), which are generally not conserved in bacteria. Conversely, in Clade B, these immune-related pathways were not enriched, but proteasomes and proteolysis pathways including eukaryotic-like E3 ubiquitin ligase gene were enriched (Figure 3C).

### Different symbiotic strategies utilizing eukaryotic-like genes

We further conducted a comparative genome analysis focusing on eukaryotic-like domains and found that all *Endozoicomonas* spp. used in this study had at least one eukaryotic-like domain (Figure 4A). The types of eukaryotic-like domains were different between Clade A1 (YOMITAN1, ONNA1, and ONNA2) and Clade B (ISHI1 and SESEKO1) *Endozoicomonas;* namely, the eukaryotic-like domain found in common between Clade A1 and Clade B was only RING-type zinc finger (RINC-type zinc-finger, Zinc-finger double domain, Zinc finger, and C3HC4 type were grouped), and the remaining six domains (SOCS box, Inhibitor of Apoptosis domain, ephrin ligand, Ets domain, MATH domain, and FANCLE C-terminal domain) were found only in either species (Figure 4A, Supplementary Table 4). We have previously reported that *Endozoicomonas* sp. ISHI1 has coral-like ephrin ligand genes [55]. In addition, there were other eukaryotic-like genes (such as the SNARE domain, Vps4 C terminal oligomerization domain-containing genes). In contrast, *Endozoicomonas* sp. YOMITAN1, ONNA1, and ONNA2, which belong to Clade A1, did not possess ephrin ligand, but had eukaryotic-like genes, including MATH/TRAF, SOCS box, an Ets-domain, and an inhibitor of apoptosis domain (Figure 4A).

**Figure 4.**
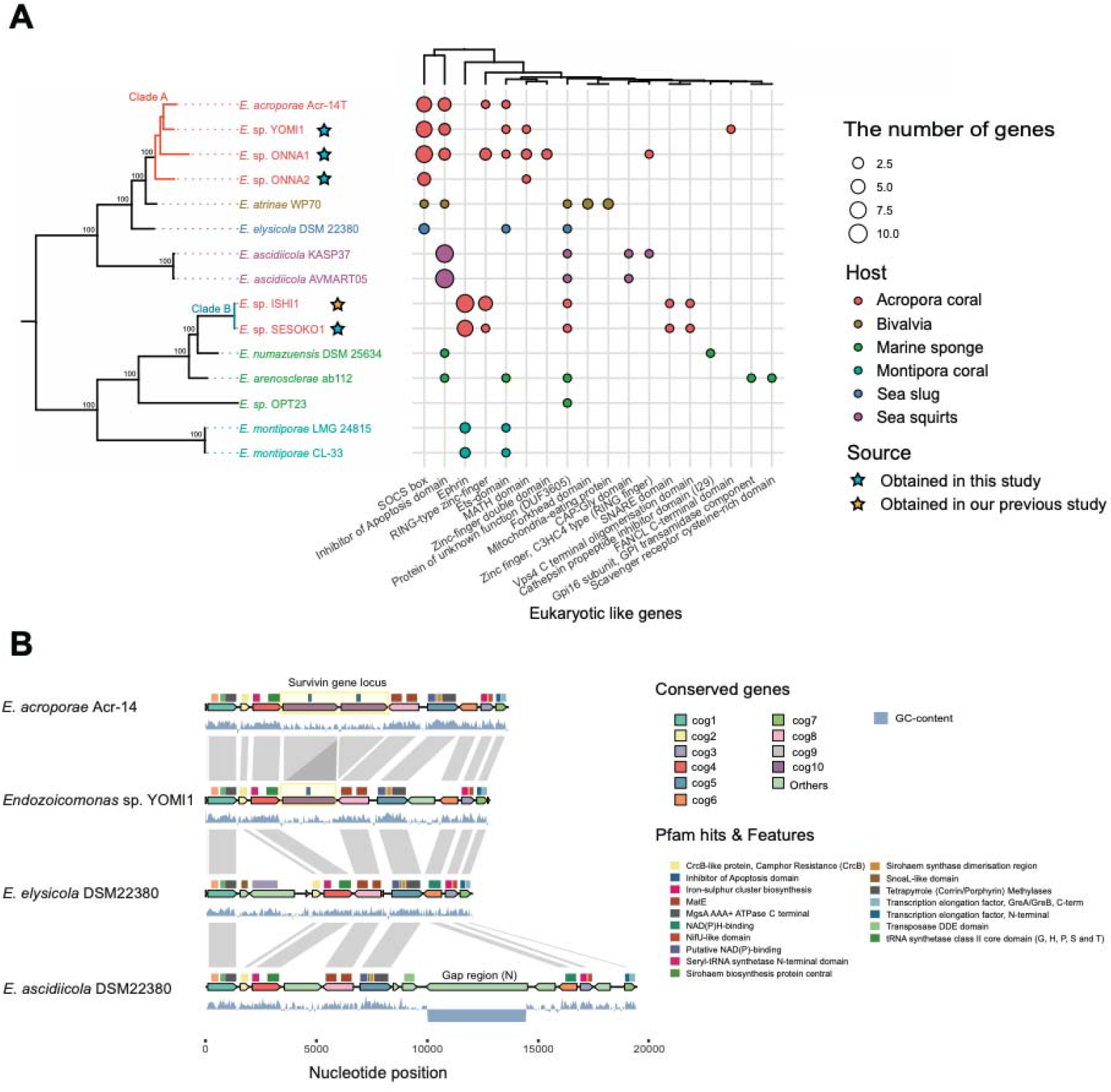
Eukaryotic-like genes distribution and horizontal gene transfer in *Endozoicomonas* A: Properties of eukaryotic-like domains predicted using EffectiveDB in *Endozoicomonas* (Z-score = 10000). The phylogenetic tree based on the single-copy marker gene is the same as that in Fig. 3A. The hierarchical clustering at the top is based on the profiles of eukaryotic-like genes. Synteny plots were created by gggenomes [71]. B: Insertion of the inhibitor of apoptosis domain-containing genes of *Endozoicomonas* sp. YOMITAN1 and *Endozoicomonas acroporae* Acr-14 into the genome. The inserted eukaryotic-like gene is marked by a yellow square. The region where the GC content of *Endozoicomonas ascidiella* DSM22380 is flat is the gap region (N).

Furthermore, we found 532 genes that did not contain eukaryotic-like domains but were highly homologous against eukaryote in *A. tenuis*-symbiotic *Endozoicomonas* (YOMI1: 125 genes, ONNA1: 134 genes, ONNA2: 47 genes, SESOKO1: 90 genes, ISHI1: 136 genes) (Supplementary Table 5). As genes closely related to host *A. tenuis* coral genes, TRAF/MATH domain-containing genes (match: 26.0-62.2%), low-density lipoprotein receptor gene (match: 30.9-67.6%), and transcription factor 15-like gene (match: 27.5-30.4%) were detected in *Endozoicomonas* Clade A1. In contrast, the genes for ephrin ligand (match: 31.3-46.0%) and RING finger and CHY zinc finger domain-containing gene (match: 46.1-64.3%) were detected in Clade B. Eukaryotic-like E3 ubiquitin-protein ligase was detected as a common gene in both clades (Supplementary Table 5). Some of these eukaryotic-like genes were inserted into the genome (eg. survivin gene, ephrin gene) (Figure 4B). Furthermore, these genes did not contain introns, despite their similarity to the intron-containing host genes (Supplementary Figure 4).

## Discussion

### The absolute abundance of *Endozoicomonas* varies greatly among coral habitats

Previous studies have focused solely on bacterial composition and failed to establish a relationship between the amount of coral symbiotic bacteria and their host conditions. To overcome this limitation, this study quantified the absolute 16S rRNA gene copy number of bacteria, and revealed the abundance of *Endozoicomonas* spp., which varied by more than 72-fold among sampling sites. Several researchers have considered that the abundance of *Endozoicomonas* is related to health and robustness of corals [56]. However, in August 2016, corals from Ishikawabaru, which had a higher abundance of *Endozoicomonas* than Sesoko Minami, showed far extensive bleaching, which was not consistent with their findings [56]. Therefore, *Endozoicomonas* in Clade B, which was abundant in Ishikawabaru, may not be related to bleaching tolerance. These results suggest that there are functional differences between *Endozoicomonas* clades and *Acropora* coral. For a deeper understanding of *Endozoicomonas* function in *Acropora* coral, the difference ofthe clade or absolute amount of *Endozoicomonas* should be considered.

### Targeted single-cell genomics enables selective accumulation of low-abundance bacterial genomes

In addition to the 16S rRNA gene-based phylogenetic analysis, we conducted singlecell whole genome sequencing of target bacteria and collected seven SAGs belonging to Clade A1 *(Endozoicomonas* sp. YOMITAN1, ONNA1, and ONNA2) and Clade B *(Endozoicomonas* sp. SESOKO1 and ISHI1). The quality of the acquired draft SAG was comparable to that of cultured strains reported previously [15], providing evidence that this method can expand genomic information of uncultured minor bacteria. It is noteworthy that our method enabled selective acquisition of draft SAGs even when the abundance ratio of target bacteria was approximately 1%, thereby overcoming the disadvantage of metagenomics. Moreover, even in the conventional single-cell genomics, the composition of sequenced bacterial taxa is highly dependent on the proportion of bacteria in the target environment because single cells are randomly sorted [57]. In contrast, our method enabled targeted single-cell genomics, showing > 50 times higher screening efficiency of target bacteria than the random sampling, thus drastically reducing the amount of reagents and labor wasted on non-target samples. Our targeted single-cell genomics can also be used to detect specific gene sequences obtained from metagenomic shotgun analysis or 16S rRNA gene sequencing by designing specific primers and probes. As a further application, specific single-cell genomics with unique characteristics will be possible by targeting the sequences of biosynthetic gene clusters and drug resistance genes. As the potential of rare bacteria to exert important functions in microbial populations is gaining increasing attention [58], our method can be a technological innovation and breakthrough for the analysis of these rare bacteria.

### Diversity of host adaptation strategies in *Endozoicomonas*

Several 16S rRNA gene amplicon and shotgun metagenomic analyses have suggested that coral-associated *Endozoicomonas* spp. may improve host coral health [56]. However, owing to the lack of reference genomes, detailed analysis of clade-specific functions in *Endozoicomonas* in the same coral species has not been performed. Therefore, we used our SAG data and compared the gene functions detected in each clade of *Endozoicomonas.* Comparative genome analysis revealed that the orthogroups of *Endozoicomonas* were conserved in each clade. Notably, even when the different clades of *Endozoicomonas* coexisted with *A. tenuis,* the shared orthogroups were only the core gene between clade A1 *(Endozoicomonas* sp. YOMI1, ONNA1, and ONNA2) and Clade B *(Endozoicomonas* sp. ISHI1 and SESOKO1). These results suggest that *Endozoicomonas* independently adapts to its host during the process of evolution.

A comprehensive search for eukaryotic-like genes in *Endozoicomonas* revealed that various eukaryotic-like genes were enriched in a lineage-specific manner. Symbiotic bacteria are known to harbor eukaryotic-like domains, such as ankyrin repeat and WD40 repeat, which are involved in protein-protein interactions and are thought to be necessary for interaction with the host [59, 60]. For example, in plants and sponges, many symbiotic bacteria and pathogens intervene in host signaling pathways via eukaryotic-like genes that mimic host genes, thereby influencing host conditions [61,62]. There have been few reports on how symbiotic bacteria acquire these eukaryotic-like genes [63], but in our study, some of these genes (eg. ephrin ligand and survivin) were found to be inserted with spliced host coral genes (Supplementary Figure 4). This result provides an implication that the insertion of these genes were mediated by host mRNA.

A recent study reported that the copy number of ankyrin repeats is higher in *E. acroporae* genomes than in other marine organism-associated *Endozoicomonas* genomes [10], implying that some eukaryotic-like genes play essential roles in the symbiosis of *Endozoicomonas* with *Acropora* coral. On the basis of this idea, we hypothesized the functions and host adaptation strategies of each *Endozoicomonas* clade based on the functions of enriched eukaryotic-like domains. Recently, it was hypothesized that ephrin ligands are required for *E. montiporae* to transition the symbiotic mechanisms [14]. In our previous report, we have also confirmed a gene expansion of ephrin ligand in *Endozoicomonas* Clade B *(Endozoicomonas* sp. ISHI1 and SESOKO1) with a higher degree than that in *E. montiporae* [55]. Interestingly, this is consistent with the fact that the number of ephrin-like receptor genes is expanded in host *Acropora* coral compared with that in *Montipora* coral [64]. These results reinforce the hypothesis that *Endozoicomonas* uses ephrin ligand gene as a part of its adaptation strategy to its host coral. The consistency of ephrin ligand and host ephrin receptor gene expansion suggests the co-evolution between *Endozoicomonas* and host coral. In other words, *Endozoicomonas* have restricted their lifestyle for establishing a symbiosis within *A. tenuis.* Furthermore, *Endozoicomonas* Clade B encodes genes associated with intracellular invasion, such as SNARE domains, which contribute to the inhibition of degradation by phagosomes in *Chlamydia* and *Legionella* [65]. However, *Endozoicomonas* Clade B does not encode immune-related genes. This result is consistent with our recent finding that *Endozoicomonas* sp. ISHI1 stimulates immune-related pathways in corals through infection (Figure. 5A) [55]. In addition, genes containing the MATH/TRAF, SOCS box, and inhibitor of apoptosis domain were present in Clade A1, but not in Clade B. These genes suppress apoptosis induced by the innate immune pathway, such as the NF-κB and JAK-STAT pathways [66]. Recently, several studies reported that the innate immune pathways are activated in host corals as stress responses to bleaching [67]. In particular, the NF-κB pathway is conserved in cnidarians and has attracted much attention in the analysis of bleaching [68–70]. Therefore, we speculate that a group of eukaryotic-like genes that are significantly enriched in *Endozoicomonas* Clade A1 may affect host innate immune pathways and increase tolerance to environmental stress. This hypothesis is consistent with the fact that *Endozoicomonas* Clade A1 in Sesoko Minami was not bleached in August 2016 (Figure 1C).

**Figure 5.**
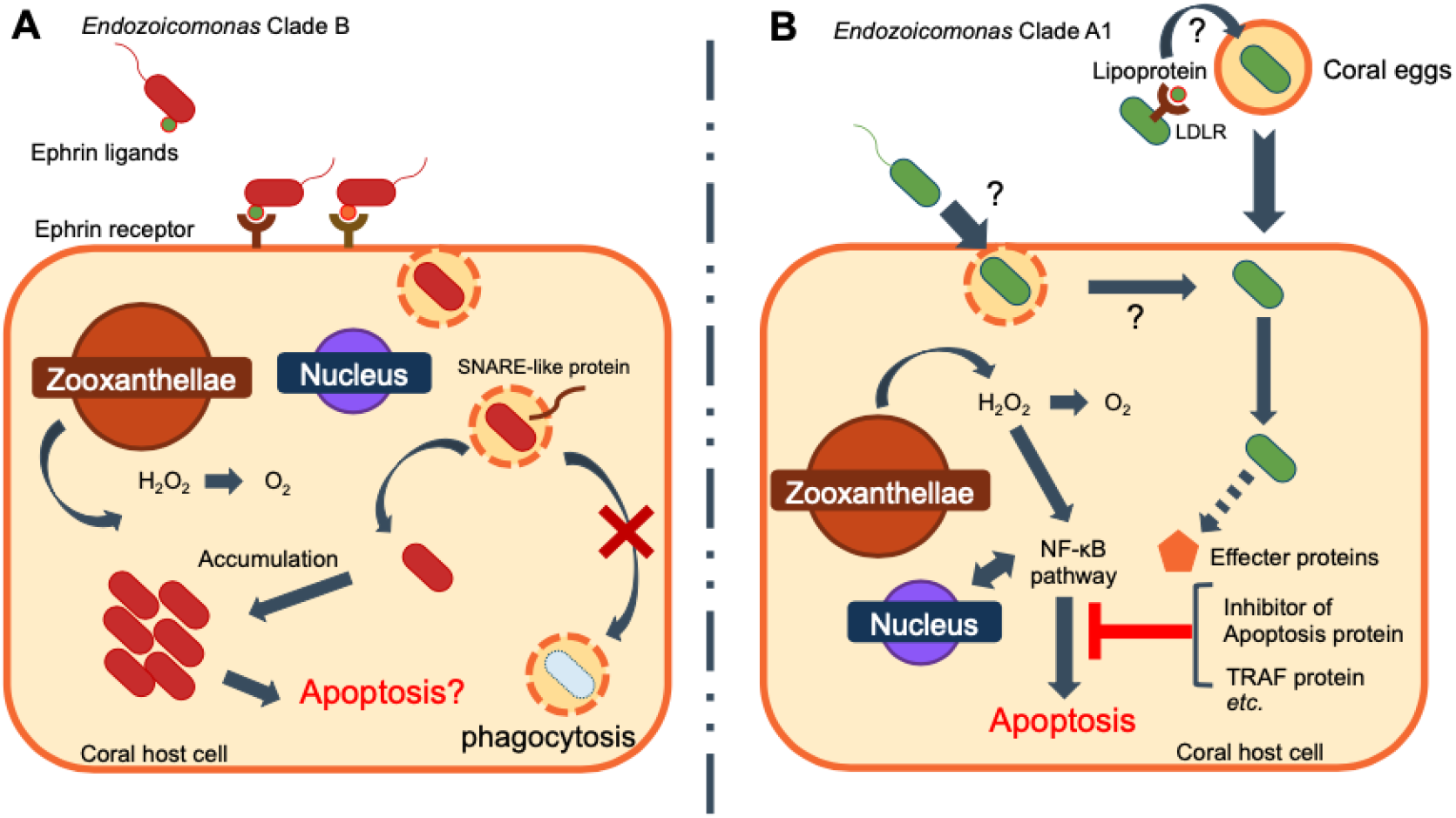
Proposed symbiotic strategy of *Endozoicomonas acroporae* in *Acropora tenuis,* as estimated by comparative genome analysis. A: Relationship between Clade B *Endozoicomonas* and *A. tenuis* corals. *Endozoicomonas* possesses many genes related to intracellular invasion, such as SNARE domain-containing genes, in addition to the eukaryotic-like gene ephrin ligand, but it does not possess any genes related to apoptosis. B: Relationships between *Endozoicomonas* Clade A1 and host *A. tenuis* corals. *Endozoicomonas* clade A1 possesses several eukaryotic-like genes involved in the suppression of apoptosis but does not have the coral-like ephrin ligand.

Homology-based analysis also revealed several eukaryotic-like genes with unknown functions in *Endozoicomonas* Clade A1 and B. These genes are phylogenetically conserved and may have some function in host adaptation and symbiosis. One example is the low-density lipoprotein receptor (LDLR) gene conserved in Clade A1. Recently, the function of LDLR during oogenesis in *A. tenuis* has been reported; it has been shown that the transcript of AtLDLR is present in the mesentery, mesenteric membrane, and mesenteric filaments surrounding an oocyte, but not in the membrane of a developing oocyte. AtLDLR can bind to major yolk lipoproteins, and may be transported into the cytoplasm of an oocyte as it develops [71]. This indicates that the LDLR-like gene of *Endozoicomonas* Clade A1 has the binding function to host egg lipoprotein. Therefore, we speculate that the LDLR like gene, which is involved in egg development, may contribute to vertical transfer of *Endozoicomonas,* as other infection-related genes such as ephrin ligand gene and SNARE gene have not been identified in Clade A1. The speculation can be verified by future experiments.

### Conclusions and perspectives

We succeeded in the targeted single-cell genomics of *Endozoicomonas* from *A. tenuis* corals with droplet microfluidics. In the single-cell genomics of environmental bacteria, it is often important to process the samples immediately after collection to obtain high-quality draft SAGs. Because the microfluidic system used in this study is portable, on-site single-cell isolation and genome amplification can be performed with standard biological laboratory equipment, which is critical for obtaining high-quality SAGs. Thus, we accomplished obtaining high-quality *Endozoicomonas* SAGs (Clade A1 and B) from the same coral species, which were proved to be phylogenetically and functionally diverse. Comparative genome analysis revealed the difference in the eukaryotic-like gene profiles, suggesting that the symbiotic strategy of *Endozoicomonas* in *A. tenuis* is divided into two types. Our findings suggest that even in the same bacterial genus, each bacterial lineage has different gene profiles and functions. Therefore, it is necessary to conduct strain-level genome sequencing to analyze the functions and strategies for host adaptation. Although *in vitro* studies in the field environment are required to prove this hypothesis, our targeted single-cell genomics could be a powerful tool for acquiring SAGs of uncultured bacterium to infer its function. Analysis of various coral-associated *Endozoicomonas* SAGs will clarify their host adaptation strategies in more detail and shed light on some aspects of the symbiotic mechanisms.

## Data availability

The assembled genomes of seven *Endozoicomonas* were deposited in DDBJ/ENA/GenBank under BioProject number PRJNA721663. The raw read data of 16S rRNA gene are available under BioProject number PRJNA722798.

## Authors Contribution

M.I., M.H., S.S., and H.T. designed the experiments. K.I., Y.Ni., Y.T., K.Y. and H.T. wrote all the manuscripts. M.I., Y.O., K.I., Y.Ni., H.I., K.K. and Y.Na. collected the data. Y.Ni. and M.K. conducted the single-cell genomics experiments for obtaining *E.* sp. YOMI1, SESOKO1-4, and ONNA1-2 SAGs. N.T., R.M. and H.I. conducted the ddPCR analysis. H.I. conducted CT scan analysis. K.I., M.K., R.W. and T.M performed the bioinformatics analysis. All authors have read and approved the final manuscript.

## Acknowledgement

We would like to thank Dr. Mitsuo Umezu for CT scan analysis, Naoko Okada for 16S rRNA gene sequencing, and Dr. Hisashi Anbutsu and Eisuke Iwamoto for useful suggestions. The super-computing resource was provided by the Human Genome Center (University of Tokyo). This study was supported by the Collaborative Research of Tropical Biosphere Research Center, University of the Ryukyus.

---

**Supplementary Figure 1.**
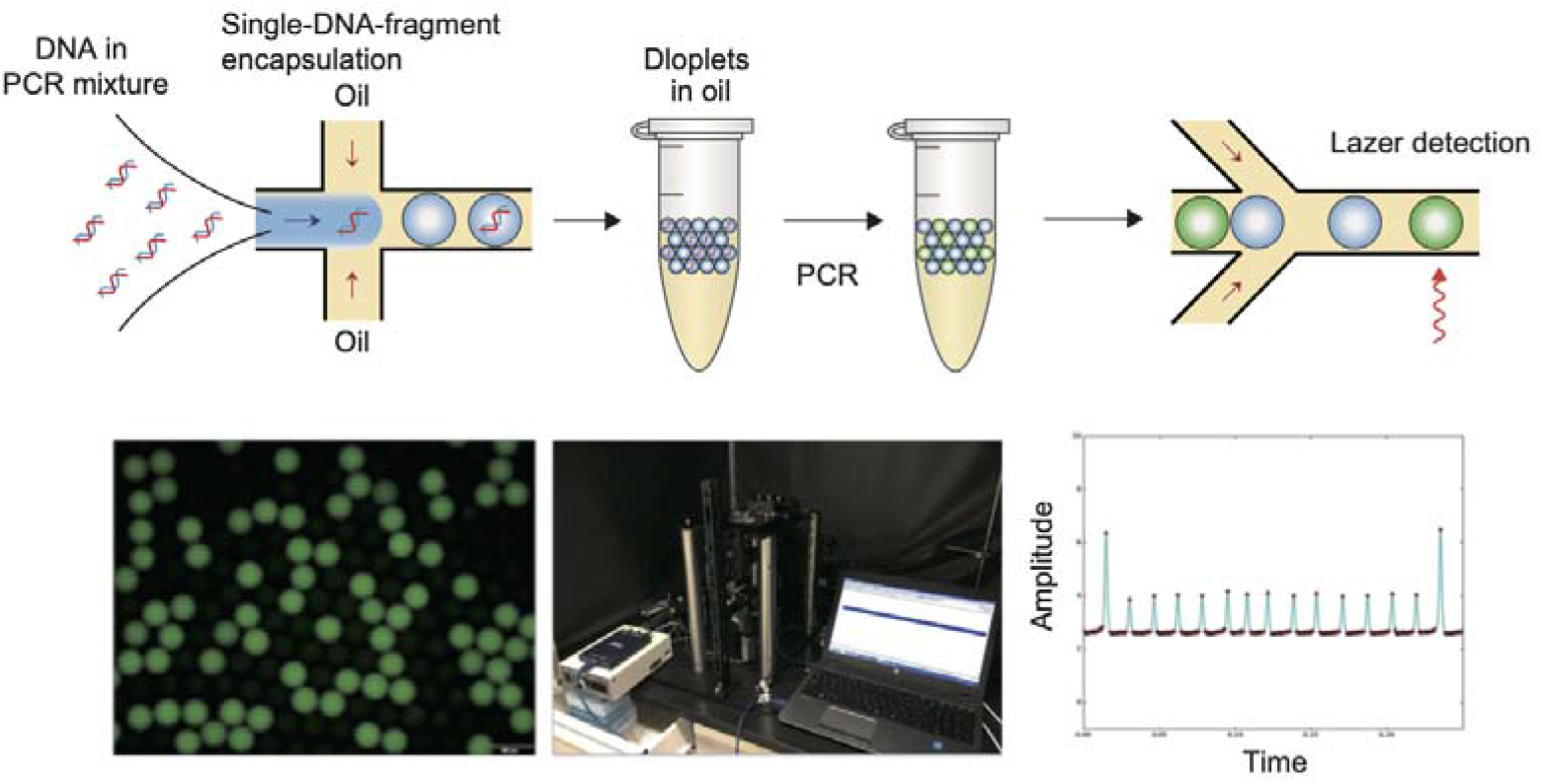
Droplet Digital PCR to quantify 16S rDNA copy number of bacteria The PCR mixture containing extracted DNA from coral branches was prepared and introduced into a microfluidic device for droplet generation. After thermal cycling, droplets were re-introduced into a microfluidic device for fluorescence detection. The number of total droplets and fluorescence-positive droplets were calculated with a custom designed optical system. The 16S rDNA copy number per 1 ng of DNA was calculated by the rate of fluorescence-positive droplets and the Poisson statistics.

**Supplementary Figure 2.**
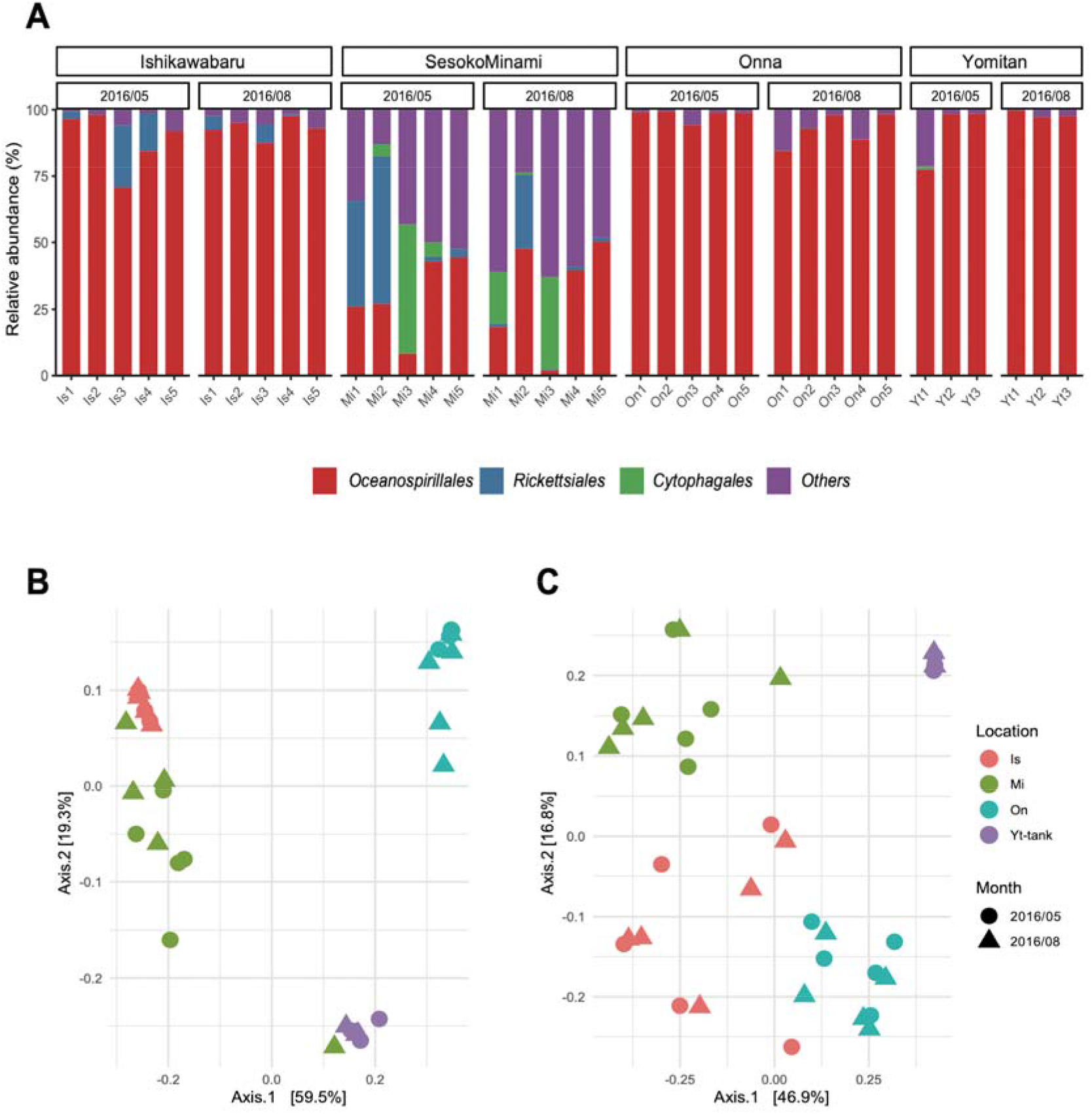
Order level barplot and PCoA plot based on UniFrac distance of *Endozoicomonas* genus in four sampling points. A: Barplot of Order level, showing the major 3 orders. B: PCoA plot of ASVs assigned to *Endozoicomonas* genus based on weighted UniFrac C: PCoA plot of ASVs assigned to *Endozoicomonas* genus based on unweighted UniFrac

**Supplementary Figure 3.**
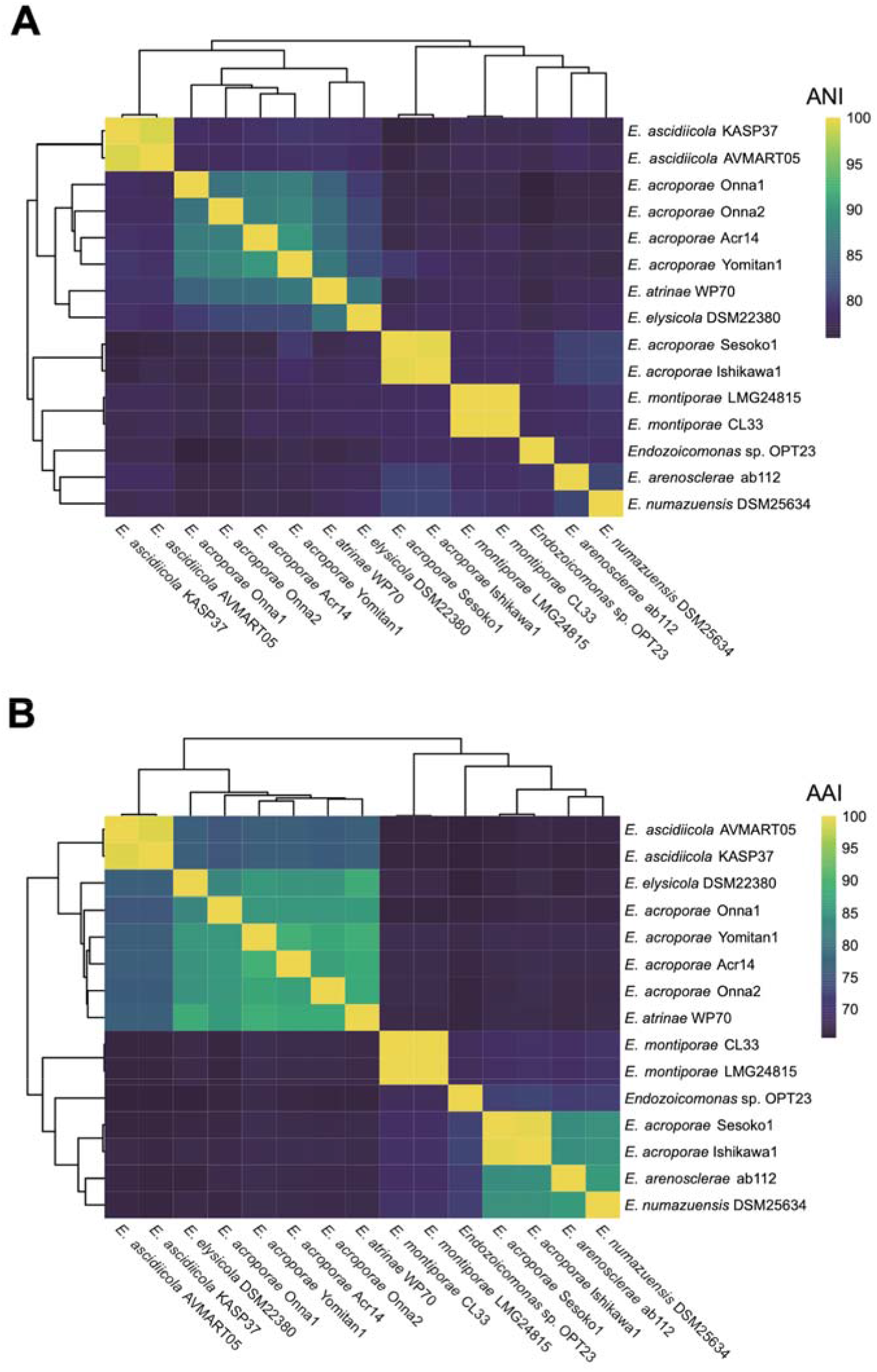
Average Nucleotide Identity (ANI) and Average Amino Acid Identity (AAI) of *Endozoicomonas* genomes. A: Average Nucleotide Identity B: Average Amino Acid Identity

**Supplementary Figure 4.**
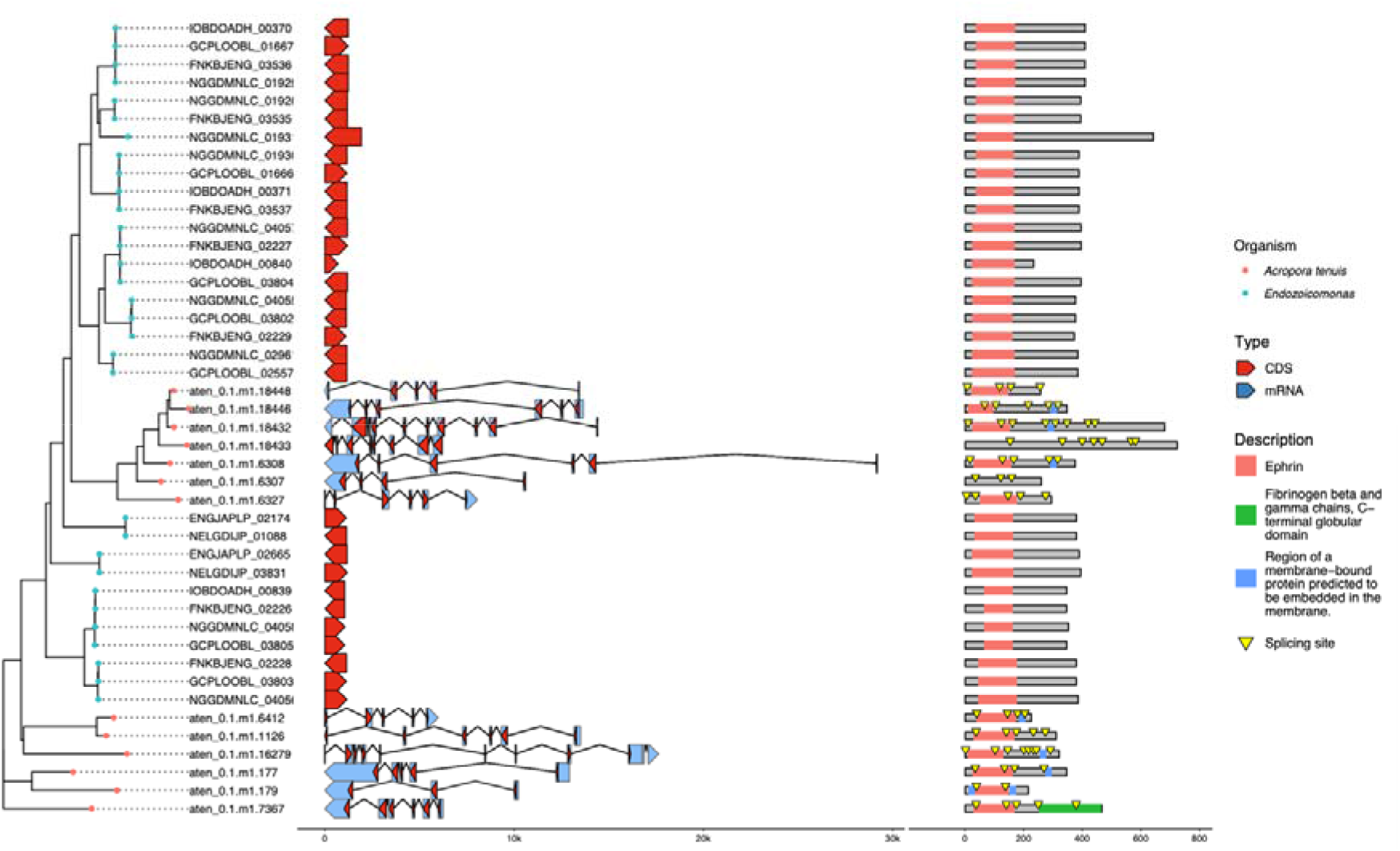
Comparison of gene structures of Coral-like ephrin genes | The left column shows the phylogenetic tree was constructed by mafft and by IQ-TREE. In the middle column, the red arrows are the coding regions for amino acids. The blue arrows are untranslated eukaryotic mRNAs. The right column shows the domain structure of the protein, and the yellow triangles indicate splicing point.

